# *Arhgap25* deficiency leads to decreased numbers of peripheral blood B cells and defective germinal center reactions

**DOI:** 10.1101/2020.03.05.965228

**Authors:** Silke E. Lindner, Colt A. Egelston, Stephanie M. Huard, Peter P. Lee, Leo D. Wang

**Affiliations:** Department of Immuno-oncology, Beckman Research Institute, City of Hope, Duarte, CA 91010, USA; Department of Pediatrics, City of Hope Medical Center, Duarte, CA 91010, USA

## Abstract

Rho family GTPases are critical for normal B cell development and function and their activity is regulated by a large and complex network of guanine nucleotide exchange factors (GEFs) and GTPase activating proteins (GAPs). However, the role of GAPs in B cell development is poorly understood. Here we show that the novel Rac-GAP ARHGAP25 is important for B cell development in mice in a CXCR4-dependent manner. We show that *Arhgap25* deficiency leads to a significant decrease in peripheral blood B cell numbers, as well as defects in mature B cell differentiation. *Arhgap25^-/-^* B cells respond to antigen stimulation *in vitro* and *in vivo* but have impaired germinal center formation and decreased IgG1 class switching. Additionally, *Arhgap25^-/-^* B cells exhibit increased chemotaxis to CXCL12. Taken together, these studies demonstrate an important role for *Arhgap*25 in peripheral B cell development and antigen response.

## INTRODUCTION

B cells are a critical part of the adaptive immune system, generating antibodies capable of recognizing a diverse array of extracellular pathogens. B cell development begins in the bone marrow, where highly regulated recombination events in B cell receptor (BCR) genes lead to the expression of an immature BCR on the cell surface (1). Immature B cells exit the bone marrow and migrate to the spleen and other secondary lymphoid organs, where they further mature as transitional B cells to become mature marginal zone (MZ), follicular (FO), or B1 B cells. Follicular B cells are the most abundant of these, located in the lymphoid follicles of the spleen and lymph nodes. It is here that the antigen- and T cell-dependent germinal center (GC) reaction occurs, which ultimately leads to the production of high-affinity antibody-secreting plasma cells and memory B cells (2–5).

Rho family GTPases are critical for B cell development and activation (6–9). Indeed, deletion of *Rac1*, *Rac2*, *RhoA*, or *Cdc42* all result in significant B cell developmental defects (6, 10–16). *In vivo*, the activity of these GTPases is moderated by a multitude of GTPase activating proteins (GAPs) and guanine nucleotide exchange factors (GEFs). GEFs like VAV and DOCK8 (17–20) have been shown to affect humoral immune responses profoundly, but the role of GAPs in B cell development and function is less-well studied. We previously showed that the Rac GAP ARHGAP25 plays an important role in hematopoietic stem and progenitor cell (HSPC) mobilization, in part by strengthening signaling through the CXCL12-CXCR4 axis to promote HSPC retention in the bone marrow niche (21). Whereas it has previously been shown that Rho family GTPases act on B cell development through actin-mediated pathways (22), here we demonstrate that *Arhgap25* plays an important role in terminal B cell differentiation and B cell activation by influencing CXCL12-CXCR4 signaling.

In particular, *Arhgap25*-deficient mice have significantly fewer peripheral blood mature B cells than wild-type controls. Additionally, *Arhgap25-*deficient B cells have defects in germinal center formation in response to *in vivo* immunization, with fewer and smaller splenic germinal centers in *Arhgap25^-/-^* mice than wild-type mice after NP-CGG injection. These defects may be attributable to increased responsiveness of *Arhgap25^-/-^* B cells to CXCL12, which may keep germinal center B cells sequestered in the dark zone (DZ)(23). We observed increased numbers of DZ B cells in *Arhgap25^-/-^* mice, as well as decreased numbers of plasma cells and diminished IgG1 secretion. Taken together, these findings indicate that *Arhgap25* plays an important role in regulating B cell chemotaxis and the germinal center reaction via the CXCL12-CXCR4 pathway.

## MATERIALS AND METHODS

### Mice

*Arhgap25^-/-^* mice (CSD28473) were obtained as cryopreserved embryos from the NIH Knockout Mouse Project (KOMP) repository, recovered using standard techniques, and housed in specific-pathogen-free barrier facilities at City of Hope. C57Bl/6N mice were purchased from Taconic Biosciences and bred under specific-pathogen-free conditions at City of Hope. Unless otherwise stated, 7–12 week old gender-matched mice were used for all experiments. Every animal was maintained and handled in accordance with City of Hope Institutional Animal Care and Use Committee (IACUC) guidelines and protocols.

### Flow Cytometry Analysis

Single cell suspensions were prepared by mechanical dissociation and then strained through a 70μm mesh. Red blood cells were lysed in RBC lysis buffer (00-4300-54, eBioscience, San Diego CA) per manufacturer’s directions. Cells were then strained through a 40μm cell strainer and then stained in phosphate-buffered saline (PBS) with 5% fetal bovine serum (FBS) in 5 ml polystyrene round-bottom tubes. Prior to antibody staining, cells were blocked with 5ng rat IgG (14131, Sigma-Aldrich, St. Louis MO). Cell surface antigens were stained with combinations of the following antibodies: CD93-FITC (AA4.1, Biolegend, San Diego CA), CD23-PE (B3B4, eBioscience, San Diego CA), IgM-PerCP-Cy5.5 (R6-60.2, BD, Franklin Lakes NJ), CD19-APC (1D3, eBioscience, San Diego CA), CD1d-Superbright 645 (1B1, eBioscience, San Diego CA), CD3e-APC (145-2C11, eBioscience, San Diego CA), CD19-BV605 (6D5, Biolegend, San Diego CA), IgD-FITC (11-26c (11–26), eBioscience, San Diego CA), CD45R/B220-BV787 (RA3-6B2, BD, Franklin Lakes NJ), CD38-FITC (90, Biolegend, San Diego CA), CD95-APC-R700 (Jo2, BD, Franklin Lakes NJ), IgG1-PE-Cy7 (RMG1-1, Biolegend, San Diego CA), IgM-BUV395 (II/41, BD, Franklin Lakes NJ), CD267-PE (eBio8F10-3, eBioscience, San Diego CA), NP-PE (N-5070-1, Biosearch Technologies, Novato CA), CD138-BV650 (281-2, BD, Franklin Lakes NJ), CD86-PE (GL-1, Biolegend, San Diego CA), CXCR4-APC (L276F12, Biolegend, San Diego CA). Cells were stained with the following viability dyes: SYTOX™ Blue Dead Cell Stain (S34857, Invitrogen, Carlsbad CA); Zombie Red (423102, Biolegend, San Diego CA); Zombie Aqua (423109, Biolegend, San Diego CA). Doublets were excluded using FSC-H/FSC-A gating. Flow cytometry analysis was performed on a BD LSRFortessa (BD, Franklin Lakes NJ) at the City of Hope Analytical Cytometry Core, and data were analyzed using FlowJo_V10 software. To determine absolute numbers of cells by flow cytometry, Precision Count Beads™ (424902, Biolegend, San Diego CA) were used. Cell count was calculated per manufacturer’s instructions.

### NP-CGG Immunization

T-cell dependent immune responses were induced by intraperitoneally injecting mice with NP-CGG (N-5055B-5, Biosearch Technologies, Novato CA), as follows: 1mg/ml NP-CGG was mixed 1:1 with freshly prepared 10% Alum (31242, Sigma-Aldrich, St. Louis MO), pH adjusted to 6.5–7.0, and washed. The precipitate was resuspended in PBS, and mice were injected with 100 μg NP-CGG. Peripheral blood was collected one week prior to NP-CGG injection and 14 days after injection. Spleens were collected for flow cytometry and histology on day 14.

### Immunofluorescence

Spleens from immunized mice were frozen in optimal cutting temperature (OCT) compound (Tissue-Tek cryomold and OCT gel compound, Sakura Finetek USA, Torrance CA) and cryosectioned in 5μm slices. Cryosections were fixed in cold acetone for 10 min at −20°C, washed with PBS, and then solubilized in 0.5% Tween-20 in PBS. After further washing in PBS, endogenous biotin and streptavidin binding sites were blocked according to manufacturer’s instructions (SP-2002, Vector Laboratories, Burlingame CA). For germinal center detection, sections were sequentially incubated with the following primary antibodies for 30 min at RT: CD3-FITC (17A2, eBioscience, San Diego CA, 1:100); CD45R/B220 (RA3-6B2, BD, Franklin Lakes NJ, 1:200); peanut agglutinin-biotin (B-1075, Vector Laboratories, Burlingame CA, 1:100). Slides were washed with 0.1% Tween-20 in PBS, and then incubated with secondary antibodies for 30 min at room temperature (AF555-goat anti-rat IgG (Invitrogen, Carlsbad CA, 1:500); AF647-sAv (Invitrogen, Carlsbad CA, 1:200)). After washing, slides were mounted with Vectashield Mounting Medium (H-1000, Vector Laboratories, Burlingame CA). Images were acquired on a Zeiss LSM 700 Confocal Microscope (magnification 200x) and pictures processed using Zen (Zeiss, Oberkochen Germany) and QuPath software.

### Luminex Assays

To assess immunoglobulin titers of NP-CGG immunized mice, serum was collected one week prior to injection and 14 days after NP-CGG treatment. Luminex multiplex assays were performed using the mouse Ig isotyping kit (MGAMMAG-300K, Millipore, Burlington MA) and conducted according to the manufacturer’s instructions by the Analytical Pharmacology Core at City of Hope.

### Chemotaxis assay

B cell migration was assessed using a 24-well, 5μm pore size transwell chamber (Corning Costar, Corning NY). Briefly, B cells were purified with CD43 micro beads (130-049-801, Miltenyi Biotec, Bergisch Gladbach Germany), and 1×10^6^ cells were placed in the top chamber of a transwell plate in 100μL of volume. 600μL of media with or without CXCL12 (100ng/ml, #250-20A, PeproTech, Rocky Hill NJ) was added to the bottom chamber. After 3h at 37°C, cells were recovered from the lower chamber, stained for cell surface makers, and analyzed by flow cytometry. As a control, 1×10^6^ cells were plated directly into the bottom chamber. The number of migrated cells was evaluated using Precision Count Beads (424902, Biolegend, San Diego CA) by flow cytometry.

### *In vitro* B cell activation

Splenic B cells of both genotypes were CD43-depleted using magnetic beads (130-049-801, Miltenyi Biotec, Bergisch Gladbach Germany) and taken in culture in the presence of 1μg/ml anti-CD40 antibody (102901, Biolegend, San Diego CA), 25ng/ml IL-4 (404-ML-025, R&D Systems, Minneapolis MN) and 25ng/ml IL-21(210-21, PeproTech, Rocky Hill NJ) for 4 days. Before culture, cells were labelled with 10μM Cell Proliferation Dye eFluor450 (65-0842-85, eBioscience, San Diego CA) according to manufacturer’s protocol. Cell division was monitored by flow cytometry and the fraction of dividing plasmablasts stained with CD138.

### Quantification and statistical analysis

All experiments described in this study were performed at least two independent times. Data were plotted and statistically analyzed using GraphPad Prism (GraphPad Software, San Diego CA). For comparisons between groups, p values were generated using a two-tailed Student t-test. Error bars represent standard deviation (SD).

## RESULTS

### *Arhgap25* deficiency results in decreased numbers of peripheral blood B cells

We previously identified *Arhgap25* as an important mediator of hematopoietic stem and progenitor cell mobilization and trafficking (21). Given the role of *Arhgap25* in Rac activation and cytoskeletal rearrangement (24, 25), and the importance of Rac activation and cytoskeletal rearrangement in B cell development and function, we reasoned that *Arhgap25* may play a role in B cell biology. We therefore investigated B cell development in 6-10 week old *Arhgap25^-/-^* mice. Interestingly, we found that *Arhgap25*-deficient animals possess significantly fewer peripheral blood leukocytes than wild-type (WT) animals (Figure 1A), and that this is accounted for by specific deficits in lymphocyte numbers (Figure 1B, C) without defects in the granulocyte or monocyte populations (data not shown). Within the lymphocyte compartment, the deficits are most pronounced in the B lymphocyte compartment (Figure 1C), and in particular among mature B cells (B220^+^IgM^+^IgD^+^, Figure 1D), indicating that there are fewer immature and mature B cells in these animals as a result of *Arhgap25* deficiency. Platelet and red blood cell numbers were unaffected by *Arhgap25* deficiency (Supplemental Fig. 1A, B).

**Figure 1.**
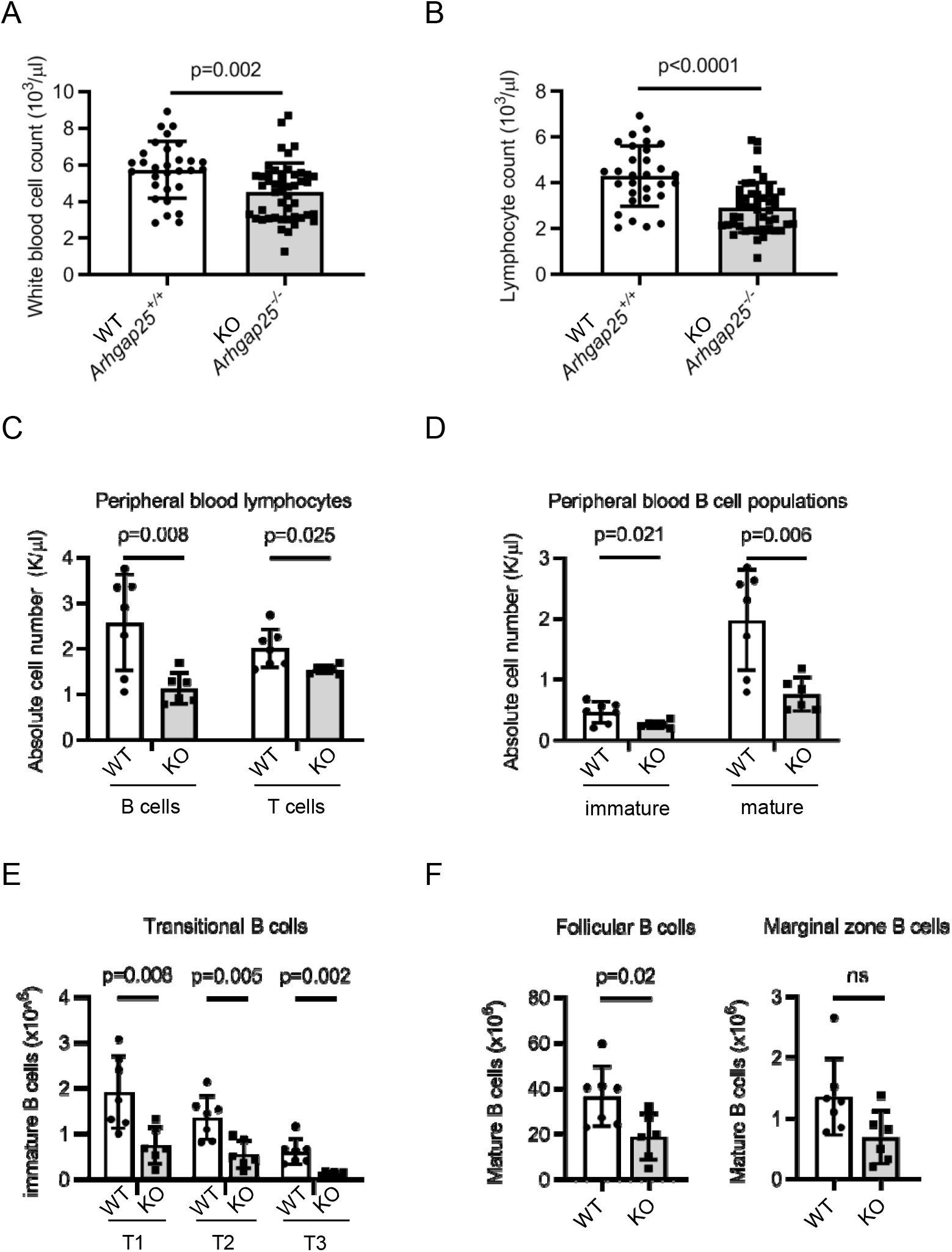
*Arhgap25* deletion leads to lower B cell numbers in peripheral blood and spleen. *Arhgap25^-/-^* (KO) mice have significantly fewer peripheral blood white blood cells (**A**) (4.54±1.56×10^3^/μl vs. 5.74±1.55×10^3^/μl, p=0.002) and fewer lymphocytes (**B**) (2.93±1.1×10^3^/μl vs. 4.3±1.32×10^3^/μl, p<0.0001) than wild-type (WT, *Arhgap25^+/+^*) mice at 6–8 weeks of age (*n*=47 KO vs. *n*=30 WT). *Arhgap25*^-/-^ mice also have fewer B (*p*=0.08) and T (*p*=0.025) lymphocytes (**C**) than WT mice, as well as fewer immature (IgM^+^IgD^-^, *p*=0.021) and mature (IgM^+^IgD^+^, *p*=0.006) B cells (**D**)(*n*=7 KO and *n*=6 WT). In the spleen, *Arhgap25^-/-^* mice have fewer transitional B cells (**E**) (T1, IgM^+^CD23^-^; T2, IgM^+^CD23^+^; T3 IgM^-^CD23^+^) and mature MZ (CD1d^hi^IgM^hi^) and FO (CD1d^+^IgM^+^) B cells (**F**) than WT mice (*n*=7 KO and *n*=6 WT). Each symbol represents one mouse; data shown as mean ± SD. *p* values calculated using two-tailed *t* test.

### *Arhgap25* deficiency affects splenic B cell development

We next investigated whether the diminution in peripheral blood B cell numbers resulted from defects in B lymphopoiesis. In the bone marrow, we found no differences in B cell developmental Hardy fractions (1) in *Arhgap25^-/-^* as compared to WT animals including the immature IgM^+^ stage (Fraction E), which is the stage at which immature B cells leave the bone marrow (Supplemental Fig. 1C, D). Thus, differences in peripheral blood B cell numbers do not appear to result from defects in bone marrow B lymphopoiesis. Immature B cells leave the bone marrow, transit through the blood, and enter the spleen as transitional (T1) B cells, where they undergo further development through the transitional T2 and T3 stages before becoming recirculating follicular (FO) or non-recirculating marginal zone (MZ) B cells (26). Consistent with our observation that *Arhgap25^-/-^* mice have fewer mature B cells in the peripheral blood, we observed decreases in the absolute numbers of transitional (T1, T2, and T3)(Figure 1E), mature follicular, and marginal zone B cells (Figure 1F) in the spleens of *Arhgap25^-/-^* mice. Taken together, these data indicate that *Arhgap25* deficiency impairs splenic B cell maturation.

### *Arhgap25*-deficient mice have impaired germinal center reactions and defective class switching

Given the defects in splenic B cell maturation in *Arhgap25^-/-^* mice, we next asked whether splenic immune responses were impaired in the absence of *Arhgap25*. 6-8 week old mice were immunized with the T-cell dependent antigen NP-CGG and their immune responses characterized 14 days later (Figure 2). Immunized *Arhgap25^-/-^* mice were observed to have fewer and smaller germinal centers (GCs) than controls (Figure 2A), as well as significant reductions in the total numbers of GC B cells (Figure 2B, C). Within the GCs of *Arhgap25^-/-^* mice, significantly more B cells were located in the dark zone (DZ) than in the light zone (LZ)(DZ:LZ ratio 4.52±1.45 in *Arhgap25^-/-^* mice vs. 2.89±0.46 in control mice, Figure 2E). Because splenic germinal centers are the major site of immunoglobulin isotype switching, we evaluated the ability of *Arhgap25^-/-^* mice to class switch. *Arhgap25^-/-^* mice had fewer IgG1^+^ NP binding GC B cells than controls (Figure 2C), as well impaired class switching to IgG1 despite equivalent serum levels of non-class-switched IgM after immunization (Figure 2C), indicating that the germinal center reaction is impaired in the absence of *Arhgap25*. Serum levels of IgM two weeks after immunization were comparable between genotypes (Figure 2D, left panel), while *Arhgap25* deficient mice revealed significantly reduced levels of IgG1 compared to control mice (Figure 2D, right panel).

**Figure 2.**
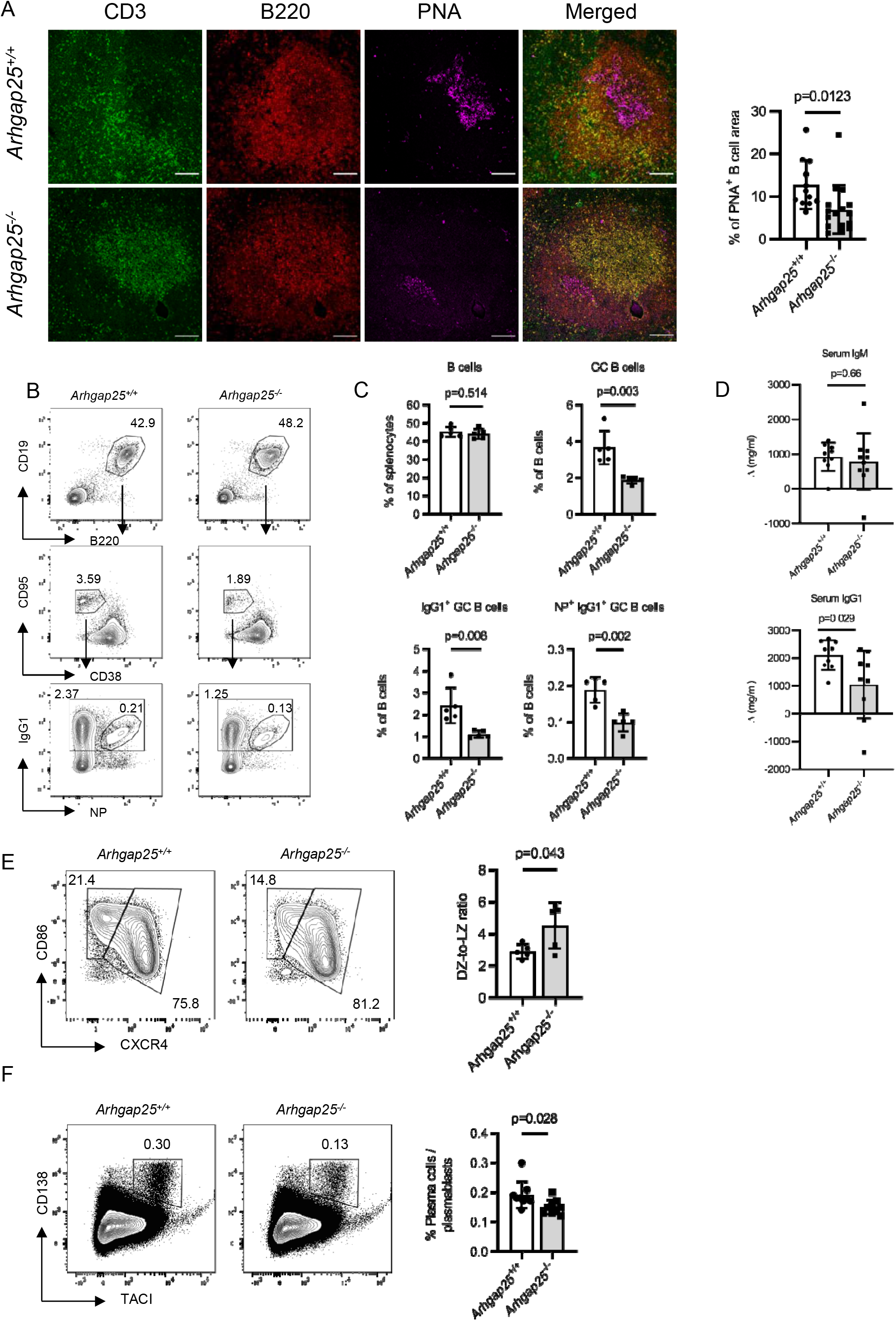
Impaired maintenance of germinal center B cells and blunted B cell response to immunization in *Arhgap25*^-/-^ mice. Mice were immunized with 4-hydroxy-3-nitrophenylacetyl chicken gamma globulin (NP-CGG) and analyzed 14 days later. (**A**) *Arhgap25^-/-^* (KO) animals have decreased germinal center (GC) formation as evidenced by a lower percentage of cells that bind peanut agglutinin (PNA). Spleen sections were stained with fluorescently labeled anti-CD3 (pseudocolored green), anti-B220 (pseudocolored red), and PNA (pseudocolored fuchsia). PNA^+^ GC area was imaged and quantified using image analysis software (n=3 mice of each genotype, scale bar=100μm). (**B**) Gating strategy for NP specific IgG1^+^ (NP^+^IgG1^+^) and total IgG1^+^ GC B cell splenocytes in (**C**). (**C**) *Arhgap25^-/-^* animals have fewer GC B cells (top right), IgG1^+^ GC B cells (bottom left), and NP^+^IgG1^+^ GC B cells (bottom right) than WT (*Arhgap25^+/+^*) animals (*n*=5 animals of each genotype). (**D**) Serum from unchallenged mice and mice 14 days post immunization was evaluated for IgG1 and IgM concentration by Luminex assay. *Arhgap25^-/-^* mice have a blunted IgG1 class switch response as compared to WT mice (bottom), whereas IgM production was equivalent before and after immunization (top). *n*=9 WT and *n*=10 KO mice, representing 3 independent experiments with 3-4 mice per genotype per experiment. (**E**) Representative contour plots (left) show gating for CXCR4^high^CD86^low^ dark zone (DZ) and CXCR4^low^CD86^high^ light zone (LZ) fractions of total GC B cells (left panel). *Arhgap25^-/-^* animals have higher DZ:LZ ratios than WT animals (right, *p*=0.043, *n*=5 mice per genotype, representative of 2 independent experiments). (**F**) Representative contour plots of TACI^+^CD138^+^ plasmablasts/plasma cells within splenic B cells 14 days after NP-CGG immunization (left panel). *Arhgap25^-/-^* animals have fewer plasmablasts than WT animals (right, *p*=0.028, *n*=5 mice per genotype, representative of 4 independent experiments).

### *Arhgap25* deficiency impairs plasma cell differentiation

Antibody production requires an efficient germinal center response and successful isotype switching, but is ultimately driven by plasma cell (PC) differentiation. We therefore evaluated the effect of *Arhgap25* deletion on PC production, and found that *Arhgap25^-/-^* mice had modestly but significantly reduced plasma cell numbers after immunization (Figure 2D). Of note, when we stimulated naive B cells *in vitro* with CD40, IL-4, and IL-21 to stimulate PC differentiation, we observed severe impairment in PC differentiation in the absence of *Arhgap25* (Supplemental Fig. 2A), indicating that the defect in PC numbers was not simply due to decreased numbers of FO B cell PC precursors. Thus, *Arhgap25* deficiency appears to impact many stages of splenic B cell development.

### *Arhgap25*-deficient B cells demonstrate increased chemotaxis to CXCL12

The CXCR4-CXCL12 interaction is of central importance in the GC reaction (23, 27, 28), and we have previously shown that ARHGAP25 regulates the strength of response to CXCR4 signaling in hematopoietic stem and progenitor cells (21). Furthermore, CXCL12 is highly expressed in the GC DZ, where we found more B cells in *Arhgap25^-/-^* mice. We therefore hypothesized that the B cell GC phenotype observed in *Arhgap25^-/-^* mice might be partly due to increased CXCR4 signaling in *Arhgap25^-/-^* B cells. To test this, we evaluated the ability of *Arhgap25^-/-^* B cells to migrate across a CXCL12 gradient; indeed, *Arhgap25^-/-^* B cells migrated at higher percentages than wild type B cells in the absence of alterations in CXCR4 expression, indicating increased responsiveness to CXCL12 (Figure 3).

**Figure 3.**
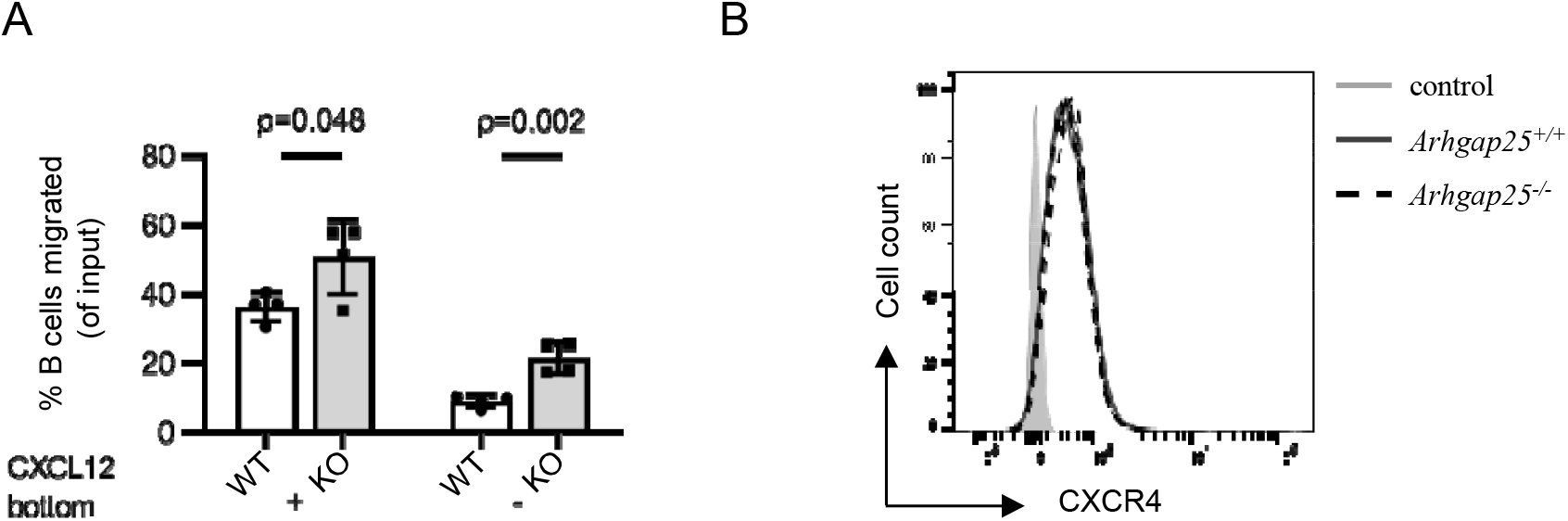
*Arhgap25*-deficient B cells accumulate in the DZ due to increased chemotaxis towards CXCL12. (**A**) *Arhgap25^-/-^* (KO) B cells migrate more effectively towards CXCL12 than do *Arhgap25^+/+^* (WT) B cells (left), and show increased motility even in the absence of a gradient (right). Data are representative of 3 independent experiments. (**B**) WT (solid grey line, MFI 508.4±7.4) and KO (dashed black line, MFI 505±22.8) B cells express equivalent levels of CXCR4 (*n*=4 mice of each genotype).

## DISCUSSION

B cells are an essential part of the adaptive immune system, responsible for producing antibodies and providing T cell help to facilitate optimal T cell responses, and proper development and differentiation of B lymphocytes is critical for immune homeostasis and host defense. While the role of small molecule GTPases such as Rac and Rho in B cell development and activation is well known, the role of Rac-GAPs is less well understood. We define the function of the novel Rac-GAP ARHGAP25 in B cell development in mice, demonstrating that mice lacking *Arhgap25* have significant defects in post-bone marrow development. Specifically, *Arhgap25^-/-^* mice have fewer splenic transitional, follicular, and marginal zone B cells as well as fewer peripheral blood B cells. They have impaired germinal center formation, isotype switching, and plasmablast differentiation. They also have increased responsiveness to CXCL12, which may cause them to accumulate in the germinal center dark zone, impairing class switch and further development. These findings are reminiscent of the human WHIM (warts, hypogammaglobulinemia, infections, and myelokathexis) syndrome, an autosomal dominant condition caused by an activating CXCR4 mutation that impairs receptor downregulation and increases CXCL12-CXCR4 signaling. The WHIM mutation results in much greater hyperresponsiveness to CXCL12 than *Arhgap25* deletion, so it is not surprising that the WHIM phenotype is much more extreme than the phenotype we report. In humans, WHIM is characterized by sequestration of neutrophils in the bone marrow (myelokathexis), as well as lymphopenia, variable hypogammaglobulinemia, and an inability to mount appropriate antibody responses (29). Mice bearing the WHIM mutation also have profound lymphopenia, impaired B and T cell development, disorganized secondary lymphoid architecture, and defects in plasma cell differentiation. However, WHIM mice do not completely phenocopy humans with the syndrome; in particular, they have normal levels of IgG (30–32). This is somewhat surprising, given that the mutation in WHIM mice impairs CXCR4 desensitization and downregulation, resulting in subsequent failure to maintain antigen responses including at the plasma cell level (30). Nonetheless, these findings support the conclusion that the B lymphopoietic defects seen in *Arhgap25^-/-^* mice are predominantly due to the effects of ARHGAP25 on CXCR4 signaling.

## ACKNOWLEDGMENTS

The authors wish to thank Maria Montoya-Arteaga and Robin Rodriguez for animal care; Verena Labi, Teresa Sadras, Martha Salas and Florian Bock for helpful discussion and suggestions; the City of Hope Analytical Cytometry Core, Analytical Pharmacology Core, Light Microscopy Core, and Pathology Core; Martha Gomez-Knight for administrative assistance; and present and former lab members.

