## Supplemental Figures for "*Arhgap25* deficiency leads to decreased numbers of peripheral blood B cells and defective germinal center reactions"

Supplemental Fig.1

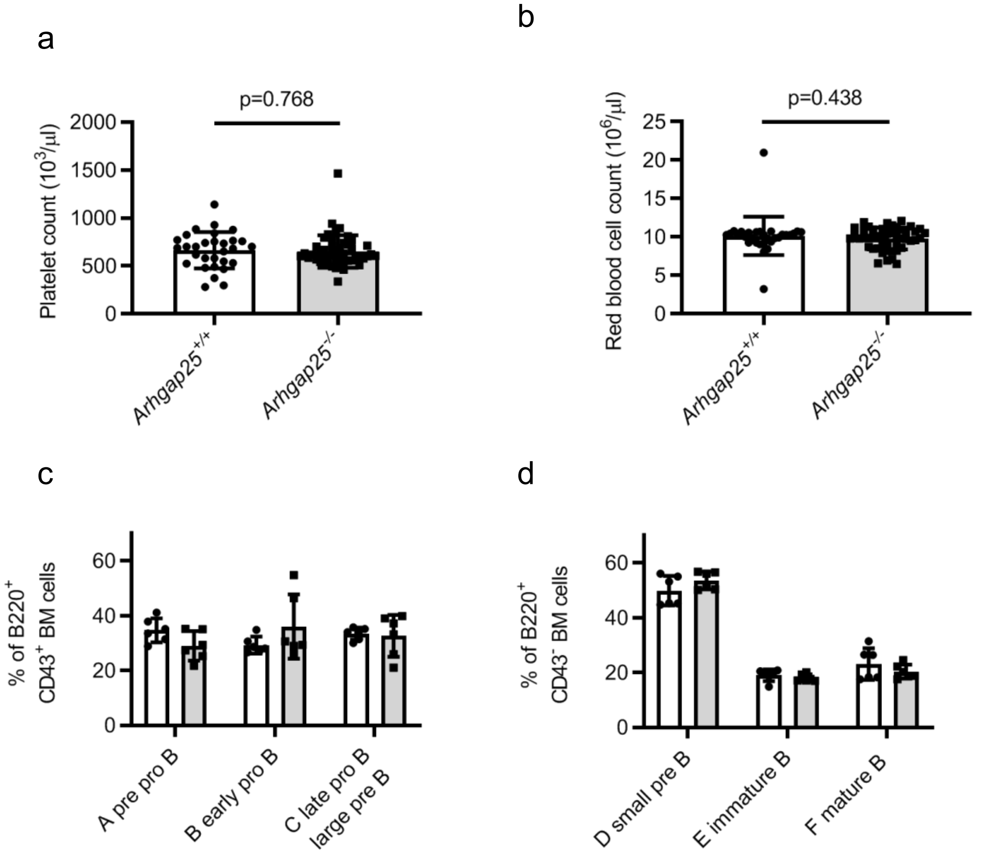


**Supplemental Figure 1.** *Arhgap25*$^{-/-}$ and WT mice have equivalent numbers of peripheral blood platelets (**A**)(652.1 ±170.8 vs. 664.4 ±189.4 K/ µL) and red blood cells (**B**) (9.8 ±1.4 vs. 10.15 ±2.5 M/µL), *n*=47 k.o. vs. *n*=30 w.t.. (**C, D**) Bone marrow B cell development is equivalent between *Arhgap25*$^{-/-}$ (grey bars) and WT (white bars) mice. (**C**) B220$^{+}$CD43$^{+}$ BM cells include Fraction A (HSA$^{-}$BP-1$^{-}$), Fraction B (HAS$^{+}$BP-1$^{-}$), and Fraction C (HAS$^{+}$BP-1$^{+}$), whereas B220$^{+}$CD43$^{-}$ BM cells (**D**) incorporate Fraction D (B220$^{dim}$IgM$^{-}$), Fraction E (B220$^{dim}$IgM$^{+}$), and Fraction F (B220$^{bright}$IgM$^{+}$). Shown are representative results from 3 independent experiments; this experiment included *n*=9 WT and *n*=10 KO mice. Each symbol represents an individual mouse. Mean ± SD are indicated; *p* values were calculated using two-tailed *t* test.

Supplemental Fig.2

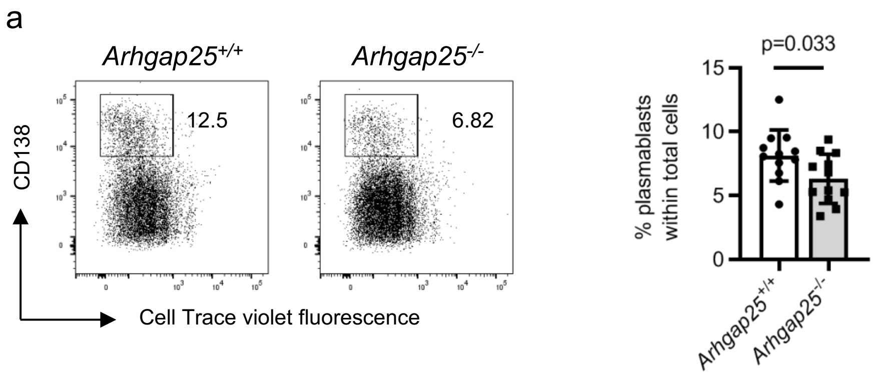


**Supplemental Figure 2.** (**A**) *Arhgap25*$^{-/-}$ B cells show defective plasma cell differentiation *in vitro*. Naïve splenic B cells (CD43$^{-}$) of *Arhgap25*$^{+/+}$ (WT) and *Arhgap25*$^{-/-}$ (KO) mice were labeled with the proliferation dye eF450 and cultured in the presence of CD40/IL-4/IL-21. On day 4, cell division and CD138 surface expression were evaluated by flow cytometry. Representative dot plots are shown at left. Right panel shows that *Arhgap25*$^{-/-}$ B cells proliferate equivalently but differentiate poorly into plasma cells; *n*=7 KO mice and 12 WT mice. Graph is representative of four independent experiments; each symbol represents an individual mouse. Mean ± SD are indicated; *p* values were calculated using two-tailed *t* test.
